# Structural basis for ligand modulation of the CCR2 conformational landscape

**DOI:** 10.1101/392068

**Authors:** Bryn C. Taylor, Christopher T. Lee, Rommie E. Amaro

## Abstract

CC Chemokine Receptor 2 (CCR2) is a part of the chemokine receptor family, an important class of therapeutic targets. These class A G-protein coupled receptors (GPCRs) are involved in mammalian signaling pathways and control cell migration toward endogenous CC chemokine ligands. Chemokine receptors and their associated ligands are involved in a wide range of diseases and thus have become important drug targets. Of particular interest is CCR2, which has been implicated in cancer, autoimmunity driven type-1 diabetes, diabetic nephropathy, multiple sclerosis, asthma, atherosclerosis, neuropathic pain, and rheumatoid arthritis. Although promising, CCR2 antagonists have been largely unsuccessful to date. Here, we investigate the effect of an orthosteric and an allosteric antagonist on CCR2 dynamics by coupling long timescale molecular dynamics simulations with Markov-state model theory. We find that the antagonists shift CCR2 into several stable inactive conformations that are distinct from the crystal structure conformation, and that they disrupt a continuous internal water and sodium ion pathway preventing transitions to an active-like state. Several of these stable conformations contain a putative drug binding pocket that may be amenable to targeting with another small molecule antagonist. In the absence of antagonists, the apo dynamics reveal intermediate conformations along the activation pathway that provide insight into the basal dynamics of CCR2, and may also be useful for future drug design.

---

The CCR2 and CCL2 signaling axis is a notable therapeutic target due to its association with numerous diseases, including cancer, autoimmunity driven type-1 diabetes, diabetic nephropathy, multiple sclerosis, asthma, atherosclerosis, neuropathic pain, and rheumatoid arthritis.^1–3^ Despite much effort that has been devoted to clinical and pre-clinical trials, a successful antagonist has yet to be developed^4–10^ (http://www.clinicaltrials.gov). Prior to a full-length crystal structure of CCR2, several studies used homology modeling and docking to gain insights into the structure and dynamics of the protein and its associated ligands or small molecule drugs.^11–13^ However, recently CCR2 was crystallized for the first time,^14^ opening up new opportunities for rational drug design.

As with most GPCRs, chemokine receptors transmit signals across cell membranes by means of extracellular ligand and intracellular G-protein binding. Distinct conformational states of the receptor are necessary for chemokine/ligand binding, G-protein binding, activation, inactivation, and signal transmission.^15–17^ GPCRs are no longer considered to be simple on/off molecular switches – they are now thought to assume a wide range of conformational states, including ligand-specific states, intermediate states, and states that allow for basal (apo) signaling without ligands bound.^16,18–24^ Ligands and small molecule drugs may shift the equilibrium of the receptor’s conformational states towards particular states. Effective small molecule antagonists that inhibit CCR2 signaling, potentially by shifting the receptor equilibrium toward inactive conformational states, are desired for treatment of diseases that involve the CCR2/CCL2 axis. Key challenges are to characterize the basal dynamics of CCR2 and to understand how the current small molecule antagonist drugs modulate these dynamics. While crystal structures provide valuable snapshots of proteins and protein complexes, they lack the ability to reveal dynamics at the atomic level. Starting with the newly resolved crystal structure of CCR2 (PDB ID: 5T1A) we performed multi-microsecond all-atom explicitly solvated molecular dynamics (MD) simulations of the receptor in a lipid bilayer in unbound (apo) and dual-antagonist-bound (holo) states (Figure 1). The two antagonists were co-crystallized with CCR2: the orthosteric antagonist, BMS-681,^14^ and the allosteric antagonist, CCR2-RA-[R]. Each system was simulated in triplicate on the Anton 2 supercomputer^25^ for a total of 260 microseconds (SI Table 1).

**Figure 1.**
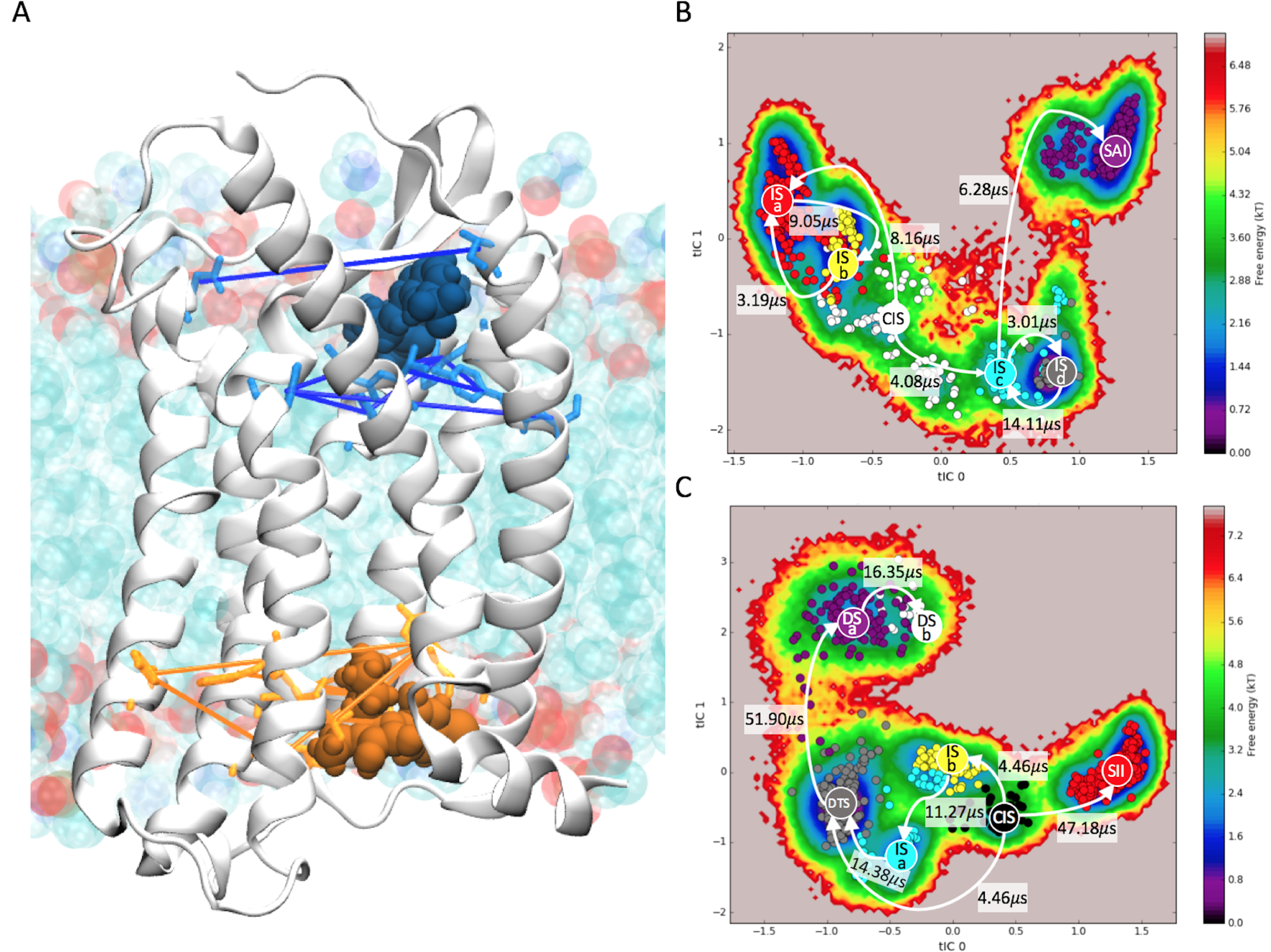
MD simulations of CCR2 in a lipid bilayer were performed on two systems: unbound (apo) and dual-antagonist-bound (holo). A) Sets of residue pairs surrounding the two ligand binding pockets were used as inputs for tICA. The protein is shown in white cartoon. Lipids are teal, red, and blue. The orthosteric ligand is shown in blue and the allosteric ligand is shown in orange. The distances between residue pairs are denoted by colored lines. The free energy and HMMs of B) apo and C) holo CCR2, projected here in the first two tICA components. Coarse-grained states are labeled and colored. Higher probability transition pathways are represented by white arrows.

While long timescale simulations are useful for analyzing sequential conformational changes, they are generally still unable to reach timescales of biological interest.^26^ One way to bridge this timescale gap is to integrate MD simulations with Markov state model theory^27–32^ (MSM, described in SI Materials and Methods). Interpreting the MD simulations with MSMs allowed us to identify key differences in the conformational ensembles and dominant slow motions of apo and holo CCR2. We find that the antagonists disrupt a continuous internal water and sodium ion pathway preventing transitions to an active-like state, and they shift CCR2 into several stable inactive conformations that are distinct from the crystal structure conformation. Three of these intermediate states reveal a putative druggable pocket that can be targeted with a small molecule inhibitor. In the absence of antagonists, we observe intermediate conformations with active-state conformational signatures that shed light on the apo dynamics of CCR2 and may also be useful for future drug design.

## Results and Discussion

### Constructing the Markov State Models

To explore the conformational landscape of both apo and holo CCR2 and identify metastable macrostates and the transitions between the macrostates, we built MSMs from MD simulation data. All simulations, barring initial equilibration runs on local resources, were performed on the Anton 2 supercomputer^25^ at the Pittsburgh Supercomputing Center. Both the apo and holo systems were simulated in triplicate, with an average of 43 microseconds per simulation (SI Table 1, Figure S9). The systems were initiated from the coordinates of the experimentally determined crystal structure.^14^ For the apo system, the co-crystallized orthosteric and allosteric antagonist ligands were removed. In each case the receptor was placed in a POPC lipid bilayer with 0.15M NaCl and solvated with TIP3P molecules.

We used time-structure independent component analysis (tICA)^33,34^ starting with all pairwise inter-residue distances to perform dimensionality reduction and identify the features and collective variables (time-structure based independent components (tICs)) that best represent the dominant slow motions in the apo and holo simulations. These features are the sets of distances between residues in the orthosteric and allosteric ligand-binding pockets (Figure 1A) [for further details on the methodology, see the SI].

One MSM was built on the apo data and a second MSM was built on the holo data. In each case, we projected the data into tICA space and clustered the trajectory frames using K-means clustering implemented in PyEMMA.^35^ After clustering, the two MSMs were built and tested. The selected models are Markovian, as indicated by implied timescale plots (Figures S1A, S2A), with a maximized number of conformational states. The Chapman-Kolmogorov test^36^ was used to test the consistency between the MSMs and the MD simulations (Figures S1B, S2B). The chosen apo MSM has a lag time of 14.4 nanoseconds and 665 clusters; and the chosen holo MSM has a lag time of 48 nanoseconds and 790 clusters. The MSMs were coarse-grained with PCCA+ to identify metastable states and their representative structures; hidden Markov models (HMMs) were used to identify transitions between those states. The apo MSM is coarse-grained into six macrostates; the holo MSM into seven macrostates (Figure 1B,C). Representative structures from each macrostate were determined by taking the centroid of the most populated cluster.

For one MD replicate of the holo system, we observed the orthosteric drug dissociate from the protein. Although there is only one dissociation event, analyzing the conformations before and after ligand dissociation gives us a first glimpse the allosteric effect of the remaining antagonist on the protein dynamics, and provides a starting point for future rounds of adaptive sampling to obtain robust dissociation statistics, which is not pursued here.

### The differential dynamics of apo and holo CCR2

The antagonists clearly affect the dynamics and kinetics of CCR2 (Figure 1B,C). Apo CCR2 transitions between each macrostate at a similar, relatively quick rate; the slowest rate of transition is 14.11 *µ*s, the fastest 3.01 *µ*s. Holo CCR2 has several transitions with rates longer than 14 *µ*s, two of which take approximately 50 *µ*s to transition to the next state; the fastest transition rate is less than 5 *µ*s. The state names are as follows: Central Intermediate State (CIS); Intermediate State (IS); Stable Active Intermediate (SAI); Stable Inactive Intermediate (SII); Dissociated State (DS); Dissociated Transition State (DTS). Holo macrostates ISb, ISa, DTS, and CIS, in the center and bottom left quadrant of Figure 1C, transition quickly between each other. The large barriers to transitioning to states DSa, DSb, and SII in the holo MSM suggest that these states are rarer and initially obstructed by the starting conformation of the protein with the antagonists bound.

The motions described by tIC 0 represent the most striking difference between apo and holo dynamics. In the apo MSM, the inter-residue distances most closely correlated with tIC 0 are all a part of the allosteric (G-protein) binding pocket, whereas in the holo MSM, the inter-residue distances most closely correlated with tIC 0 are all a part of the orthosteric (chemokine) binding pocket (Figure S4).

tIC 1 in apo CCR2 represents the flipping of TRP 98^2.60^ into the orthosteric drug binding site (Figure 2A,B, Figure S4C). In the crystal structure TRP 98^2.60^ packs with the tri-substituted cyclohexane of the orthosteric antagonist, BMS-681.^14^ Without the presence of this ligand, TRP 98^2.60^ assumes three distinct positions as it transitions from the crystal structure conformation to a position where it obstructs the chemokine binding pocket. The TRP 98^2.60^ conformation in the cluster at the neutral tIC (boxed in yellow, Figure 2A,B) most closely resembles the crystal structure conformation. The two other conformations are found at the extreme ends of tIC 1 in densely populated free energy wells. Of these two conformations, State SAI assumes the most drastic conformation and protrudes into the chemokine binding site. In the holo simulations and holo macrostates there is markedly less intrusion into the binding pocket due to the presence of the ligand.

**Figure 2.**
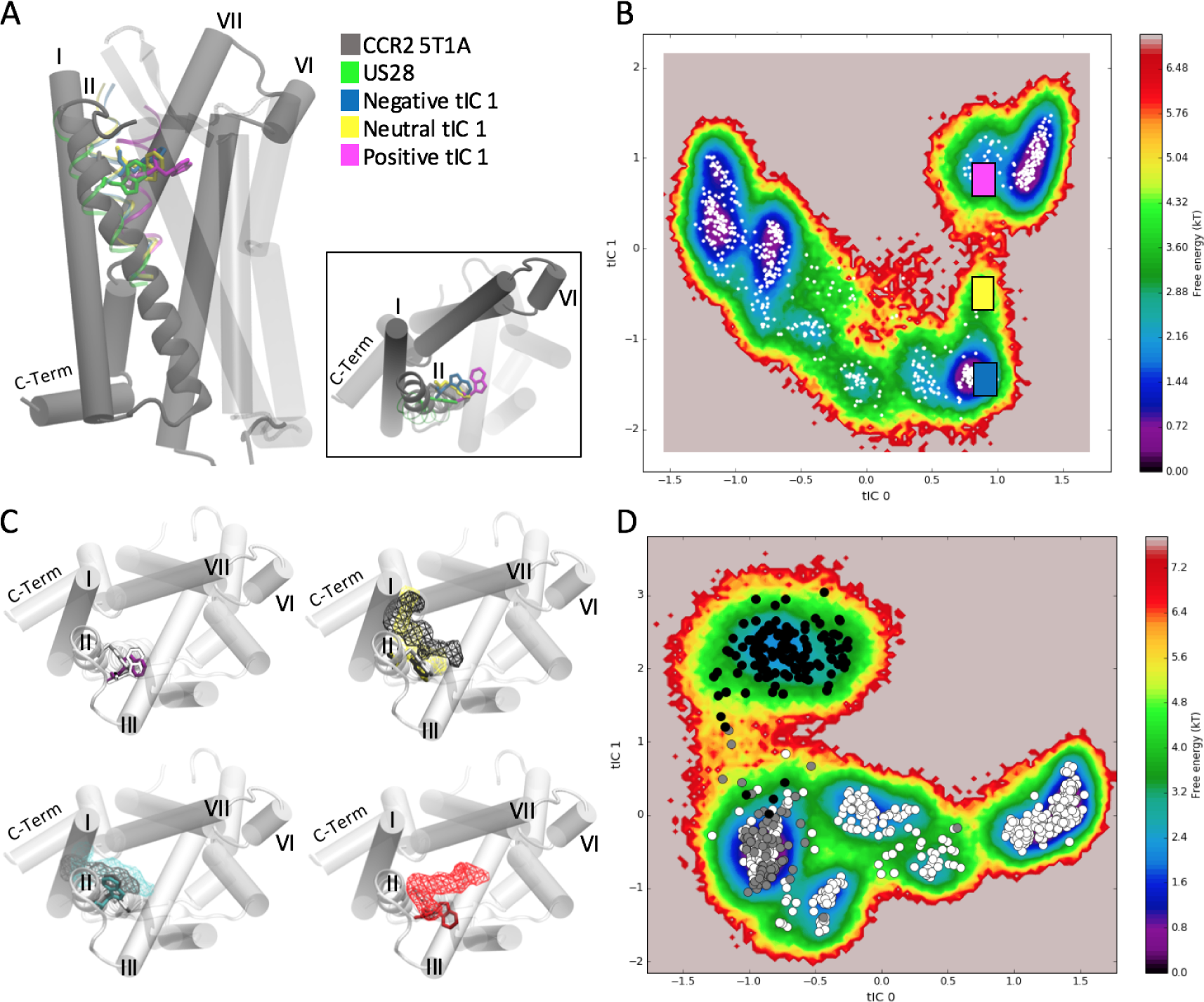
A) In apo CCR2, tIC 1 represents TRP 98^2.60^ in three distinct positions. In gray is the crystal structure; in green is the active crystal structure of US28; in blue and yellow are transitions, and in magenta is the most dramatic conformation. Each conformation is plotted on the free energy in tICA space in B). C) In holo CCR2, the positioning of the orthosteric ligand and the conformation of TRP 98^2.60^ is closely linked. Shown in light silver cartoon is CCR2 5T1A; TRP 98^2.60^ is displayed as purple in State DSa, white in State DSb, gray in State DTS, cyan in State ISa, yellow in state ISb, black in State CIS, and red in State SII. D) Holo CCR2. White circles are clusters of frames before any ligand dissociation. Grey circles are clusters of frames during the event. Black circles are clusters of frames after the event.

tIC 1 in holo CCR2 represents the concerted movement of 5 pairs of residues in the orthosteric ligand binding site during this event of the orthosteric ligand dissociation (Figure 2, Figure S4D). The separation projected in the first two tICs (Figure 2A) is clearly divided into clusters of frames that occur before (white clusters), during (grey), and after (black) dissociation. The transition rate between the pre- and post-dissociated clusters is the largest rate in either MSM: 51.90 *µ*s (Figure 1C). The residue pairs identified by tICA that contribute to tIC 1 and this ligand dissociation (Figure 2B) were confirmed by analyzing the original simulation data. The key changes are the change in distance between TYR 49^1.39^ - THR 292^7.40^, TRP 98^2.60^ - TYR 120^3.32^, SER 50^1.40^ - TYR 259^6.51^, and the chi angle of GLU 291^7.39^. Notably, four of these residues (TYR 49^1.39^, TRP 98^2.60^, TYR 120^3.32^, and THR 292^7.40^) are not only involved in binding to the co-crystallized orthosteric antagonist BMS-681, but are also critical for CCL2 binding and/or activation of CCR2.^37,38^

In the CCR2 crystal structure, there is a hydrogen bond between TYR 49^1.39^ and THR 292^7.40^. The gamma-lactam secondary exocyclic amine of the orthosteric ligand forms a hydrogen bond with the hydroxyl of THR 292^7.40^, and the carbonyl oxygen of the gamma-lactam forms a hydrogen bond with TYR 49^1.39^. During simulation, the distance between TYR 49^1.39^ and THR 292^7.40^ averages 0.4 nm until 3 *µ*s after the ligand dissociates, when it begins fluctuating between 1.3 nm to 0.3 nm (Figure S5). This suggests that the orthosteric ligand dissociation breaks the hydrogen bond between these key ligand binding residues. This motion is captured in the holo MSM: the separation of the two residues is exemplified between States ISa (pre-dissociation) and DSb (post-dissociation) in Figure S5A,B. The distance between SER 50^1.40^ and TYR 259^6.51^ is also a contributor to tIC 1, and shows the same outward movement of helix I. In Figure S5C, we see a slight decrease in distance between the residue pair, followed by the same lag time of 3 *µ*s as discussed above, and finally an increase in distance as the extracellular end of helix I bends away from the helical bundle.

The positioning of the orthosteric ligand and the conformation of TRP 98^2.60^ are closely linked (Figure 2C). After ligand dissociation, in holo States DSa and DSb (purple and white, respectively), TRP 98^2.60^ turns towards helix III, bending slightly inward toward the chemokine binding site. Prior to ligand dissociation, TRP 98^2.60^ has two distinct conformations. In the first conformation (States ISb and CIS, black and yellow), the ligand positions itself between helices I and VII, in the same conformation as the crystal structure. TRP 98^2.60^ is constrained in a downward position, pointing intracellularly, also resembling the crystal structure conformation. In the second conformation (States ISa and DTS, cyan and grey), TRP 98^2.60^ flips up and out of the binding pocket, pointing extracellularly, and the ligand moves between helices I and II. The third conformation of TRP 98^2.60^ is found in State SII (red), and is the most prominent position of the residue because it extends deeper into the chemokine binding site toward helix III. In this case, the ligand interacts with helices II, IV, and V, and there are no transitions from this state to a dissociated state.

As in the apo MSM, the absence of the orthosteric ligand causes a shift in the position of TRP 98^2.60^. In the holo simulations shown in Figure S6, the dissociation event is preceded by a doubling of the distance between TRP 98^2.60^ and TYR 120^3.32^, and 3 *µ*s after dissociation the distance returns to its previous 0.4 nm. This increase in distance may be required for the ligand to begin the process of dissociating. Another drastic change during the dissociation event is the switch of GLU 291^7.39^ from a constrained chi angle of −50 to −100 degrees to an unconstrained 150 to −150 degrees (Figure S7). After dissociation, this angle more closely resembles the conformation in all apo simulations.

Finally, the faster dominant motions (tIC 2, 3, 4) in the holo MSM consist of rearrangements in the allosteric ligand binding site, perhaps suggesting that an allosteric rearrangement must first happen in order for the orthosteric ligand to dissociate. Further evidence for this is the observed correlated motion of the downward flip of the conserved microswitch residue TYR 305^7.53^ in the G-protein binding site with the dissociation of the orthosteric ligand from the chemokine binding site, which is discussed further in the following section.

### Simulations reveal a wealth of conformational differences between apo and holo CCR2

Apo and holo CCR2 access unique intermediate states, revealing novel opportunities for rational drug design. The intermediate states of apo CCR2 have conformational signatures found in the activate states of GPCRs, suggesting that these states are on a pathway toward activation. Holo CCR2 diverges from the crystal structure to form distinct intermediate states that expose putative drug binding pockets and reveal the effect of antagonists on receptor dynamics.

We evaluate the metastable states by comparing helical conformational signatures and conserved microswitch residues, including NPxxY (Tyr 305^7.53^), DRY (Arg 138^3.50^), Tyr 222^5.58^, the allosteric and orthosteric sets of residues introduced above, and the chemokine and G-protein binding pockets to the inactive crystal structure of CCR2 that we used in this study (PDB ID 5T1A) and the active crystal structure of another class A GPCR, US28 (PDB ID: 4XT3; 30% sequence identity to PDB ID 5T1A). Signatures of an active GPCR state include 1) the inward shift of the intracellular part of helix VII toward the helical bundle 2) the outward shift of the intracellular part of helix VI in concert with helix V, 3) the upward shift and lateral movement of helix III, and 4) the rearrangements of conserved microswitches.^18^ According to these metrics, the starting crystal structure of CCR2 is in a particularly inactive conformation,^14^ whereas the crystal structure of US28 is in the active conformation.

#### 1) Apo macrostates show an active-like inward shift of the intracellular part of helix VII toward the helical bundle

An active state hallmark that is apparent in all of the apo macrostates is the conformation of helix VII (Figure 3A): the intracellular end of helix VII tilts slightly inward toward the center of the helical bundle. More prominently, the extracellular end of helix VII tilts outward, resembling the active conformation of US28. The holo macrostates show the opposite: the intracellular end of helix VII tilts slightly outward and the extracellular end of helix VII tilts inward, remaining in the crystal structure conformation.

**Figure 3.**
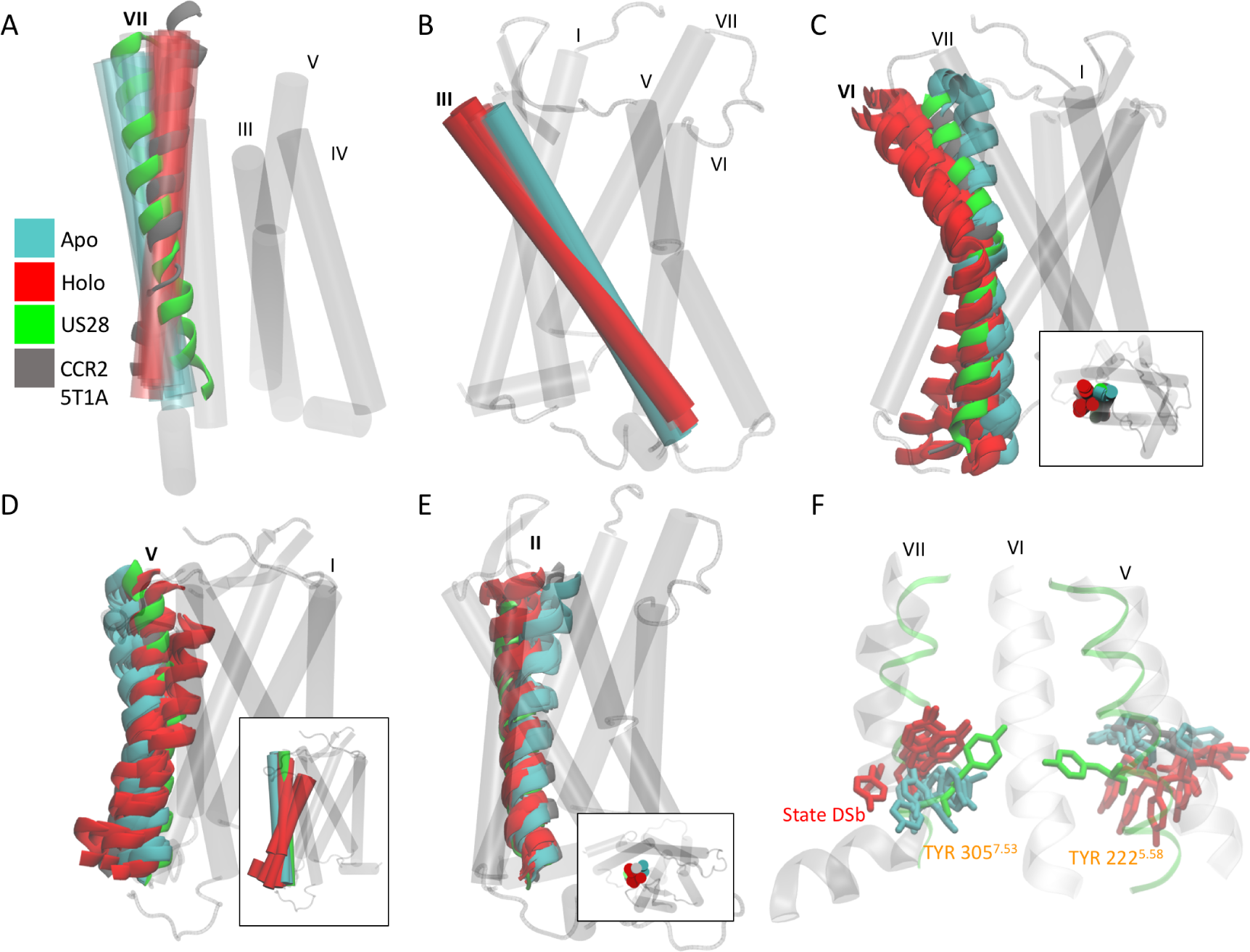
Apo and holo macrostates are compared to the active crystal structure of US28 (green, PDB ID 4XT3) and the crystal structure of CCR2 (grey, PDB ID 5T1A). All apo metastable states (teal) show A) helix VII tilting outward, B) helix III tilting upwards, C) helix VI tilting inward, D) helix V tilting outward, and E) helix II tilting inward. All holo macrostable states (red) show A) helix VII tilting inward, B) helix III tilting outward, C) the extracellular end of helix VI tilting outward, D) the extracellular end of helix V tilting inward, and E) helix II tilting outward. Insets in C), E) show the extracellular to intracellular view in cartoon. Inset in D) depicts the same view of apo and holo helix V in cartoon. F) In licorice are conserved motifs TYR 305^7.53^ and TYR 222^5.58^. All six apo metastable state assume a new conformation for TYR 305^7.53^. Six out of the seven holo metastable states have TYR 305^7.53^ in the same conformation as the equilibrated crystal structure. Post-ligand-dissociation holo state DSb assumes a new position of TYR 305^7.53^, more similar to the dominant apo conformation. Apo metastable states sample a narrower range of conformations for TYR 222^5.58^ than holo.

#### 2) Holo macrostates, not apo, show an active-like outward shift of the intracellular part of helix VI in concert with helix V

Helix V and VI in the apo macrostates are not in an active conformation. Instead, it is the holo macrostates that have the intracellular end of helix V and VI tilting outward to resemble the active conformation (Figure 3C,D), suggesting that neither apo nor holo macrostates are in an exclusively inactive or active conformational state, despite starting from a particularly inactive crystal structure.

#### 3) Apo macrostates show an active-like upward shift and lateral movement of helix III

Apo macrostates also resemble the active conformation by the slight upward shift of helix III; unlike holo macrostates, which remain in a position similar to the inactive crystal structure (Figure 3B).

#### 4) The rearrangements of conserved microswitches suggest that apo macrostates are active intermediates, and holo macrostates are inactive intermediates

##### a) NPxxY motif (Tyr 305^7.53^)

In the inactive conformation of GPCRs, Tyr 305^7.53^ points towards helices I, II, or VIII (in CCR2, it points toward II), and in the active state Tyr 305^7.53^ points toward middle axis of helical bundle.^18^ Each apo macrostate shows Tyr 305^7.53^ pointing downward into the intracellular (G-protein) binding pocket (Figure 3F). This positioning of Tyr 305^7.53^ does not match the active conformation in US28, and is also distinct from the inactive crystal structure of CCR2. In six out of the seven the holo macrostates, Tyr 305^7.53^ is stabilized in the inactive state. The holo macrostate in which Tyr 305^7.53^ is not stabilized in the inactive conformation is accessed after the orthosteric ligand dissociates (State DSb, Figure 1C); the allosteric pocket residues rearrange and Tyr 305^7.53^ assumes a conformation similar to the apo states. These concerted events may indicate allosteric cross-talk between the chemokine binding site and the G-protein binding site.

##### b) The microswitch residue Trp 256^6.48^, and the interaction of the DRY motif (Arg 138^3.50^) with Tyr 222^5.58^

Apo and holo macrostates both maintain the same chi angle of the conserved microswitch residue Trp 256^6.48^ which describes an active GPCR when it switches from gauche to trans conformation and facilitates the interaction of Tyr 222^5.58^ and Tyr 305^7.53^. The interaction of these two tyrosines and Arg 138^3.50^ also characterizes an active state GPCR.^39^ In the inactive crystal structure of CCR2, Tyr 222^5.58^ points toward lipids, sterically blocked by Phe 246^6.38^ from interaction with Arg 138^3.50^ and Tyr 305^7.53^.^14^ In apo macrostates, Tyr 222^5.58^ remains pointed toward the lipids, never swiveling around to interact with Arg 138^3.50^ or Tyr 305^7.53^ as occurs in activated GPCR states (Figure 3F). Holo macrostates actually show increased range of motion of Tyr 222^5.58^, diverging from the crystal structure to stabilize in unique intermediate conformations. The steric obstruction from Phe 246^6.38^ is alleviated in both apo and holo macrostates, as Phe 246^6.38^ swings outward and points toward the lipids. The conformations of these microswitch residues indicate that both apo and holo macrostates are sampling different intermediate conformations.

#### 5) Comparison of binding sites

The effects of the antagonists on the conformational ensemble of CCR2 are evident when comparing the binding sites of the apo and holo macrostates. In the holo macrostates there is a dramatic expansion of the extracellular (chemokine) binding site. This expansion is caused by a pronounced outward tilting of helix VI and slight outward tilting of helix II in the holo macrostates; the apo macrostates show the opposite, with a slight inward tilting of both helices VI and II (Figure 3C,E). The intracellular (G-protein) binding site also enlarges in the holo macrostates due to the outward shift of the intracellular ends of helices V and VI, but remains obstructed in all apo and holo states. In the crystal structure, this obstruction occurs by the interaction of Arg 138^3.50^ with Asp 137^3.49^ and with Thr 77^2.39^.^14^ These interactions are maintained throughout all the simulations and the chi angle of Arg 138^3.50^ remains constrained despite the slight expansion of the binding site. Interestingly, instead of disrupting these key interactions in the G-protein binding site, the outward movement of the intracellular end of helix VI and the movement of helix V toward helix VI in intermediate states DSb, DTS, and ISa in the holo MSM create a putative druggable site for novel allosteric antagonists (Figure 4). Computational solvent fragment mapping^40^ further indicates that the pocket is a binding hot spot due to its ability to bind clusters of multiple different drug-like probes (Figure 4C). The pocket can be accessed through the lipid bilayer between helices IV and V, or from the G-protein binding site, as a deeper extension of the current allosteric binding site of CCR2-RA-[R], and may be useful for rational drug design or modification of current antagonists.

**Figure 4.**
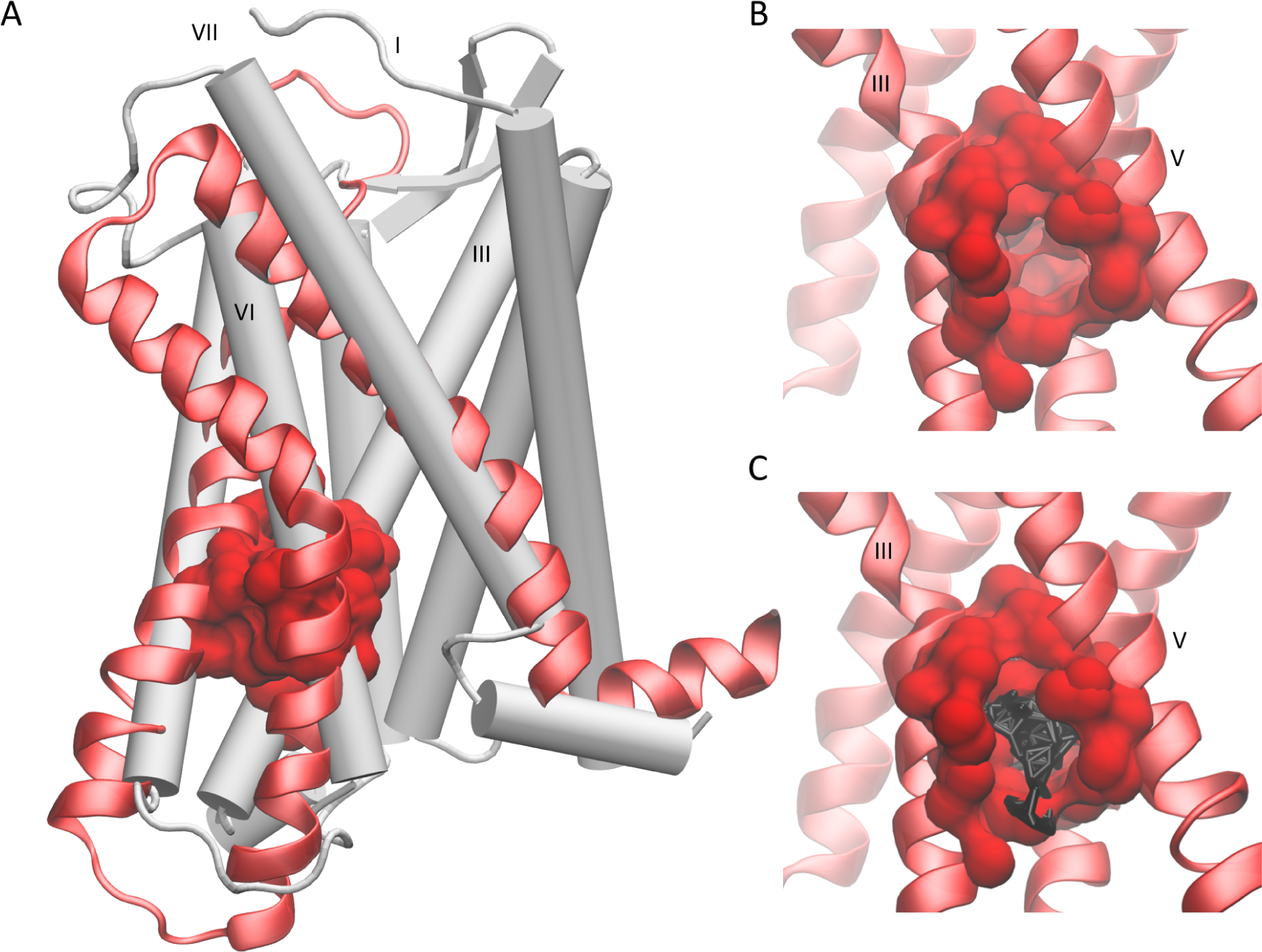
A putative allosteric drug binding pocket is revealed by three holo macrostates. A) A comparison of the CCR2 crystal structure (white cartoon) with helices V, VI, and VII (red new cartoon) of one holo macrostate. The pocket is shown in red surf. B) A closer view of the pocket from the other side of the protein, between helices III and V. C) Small organic probes used for computational fragment mapping are shown in black licorice.

Overall, holo macrostates show more helical tilting, bending, and binding site expansion, which increases the solvent-accessible surface area (SASA) when compared to the crystal structure and the apo macrostates. However, the apo simulations have greater overall increased motion and residue fluctuation, suggesting that the antagonist ligands dampen the dynamics (Figure S3).

### Antagonists disrupt internal water and sodium ion pathways

Internal water molecules are thought to be an integral part of receptor activation in GPCRs, but the exact mechanism remains unknown.^41^ Internal water molecules can strongly influence conformational changes in GPCRs by interfering with hydrogen bonding networks of the receptor’s backbone and side chains. Here, we use MD to provide detail about internal waters within CCR2 at an atomic level that is inaccessible to other experimental methods such as X-ray crystallography.

A continuous internal water pathway forms in apo CCR2 (5A). The antagonists in the holo simulations disrupt this water pathway, preventing water molecules from passing all the way through the protein core. Water occupancy per residue was calculated for apo and holo CCR2 (Figure S8A). Several of these higher water occupancy residues (e.g. Asp 36^1.26^, Ser 50^1.40^, Glu 235^6.27^, Lys 236^6.28^, Glu 310^8.48^, Lys 311^8.49^) may be exposed to more water in the apo simulations than in the holo simulations simply because the ligands have been removed and the water has access to the binding pockets. The other residues (e.g. Asp 78^2.40^, Tyr 80^2.42^, Asp 88^2.50^, Leu 92^2.54^, Ile 93^2.55^, Gly 127^3.39^, Ile 128^3.40^, Glu 291^7.39^, and Phe 312^8.50^) are located in the core of the protein, along the continuous pathway (Figure 5A).

**Figure 5.**
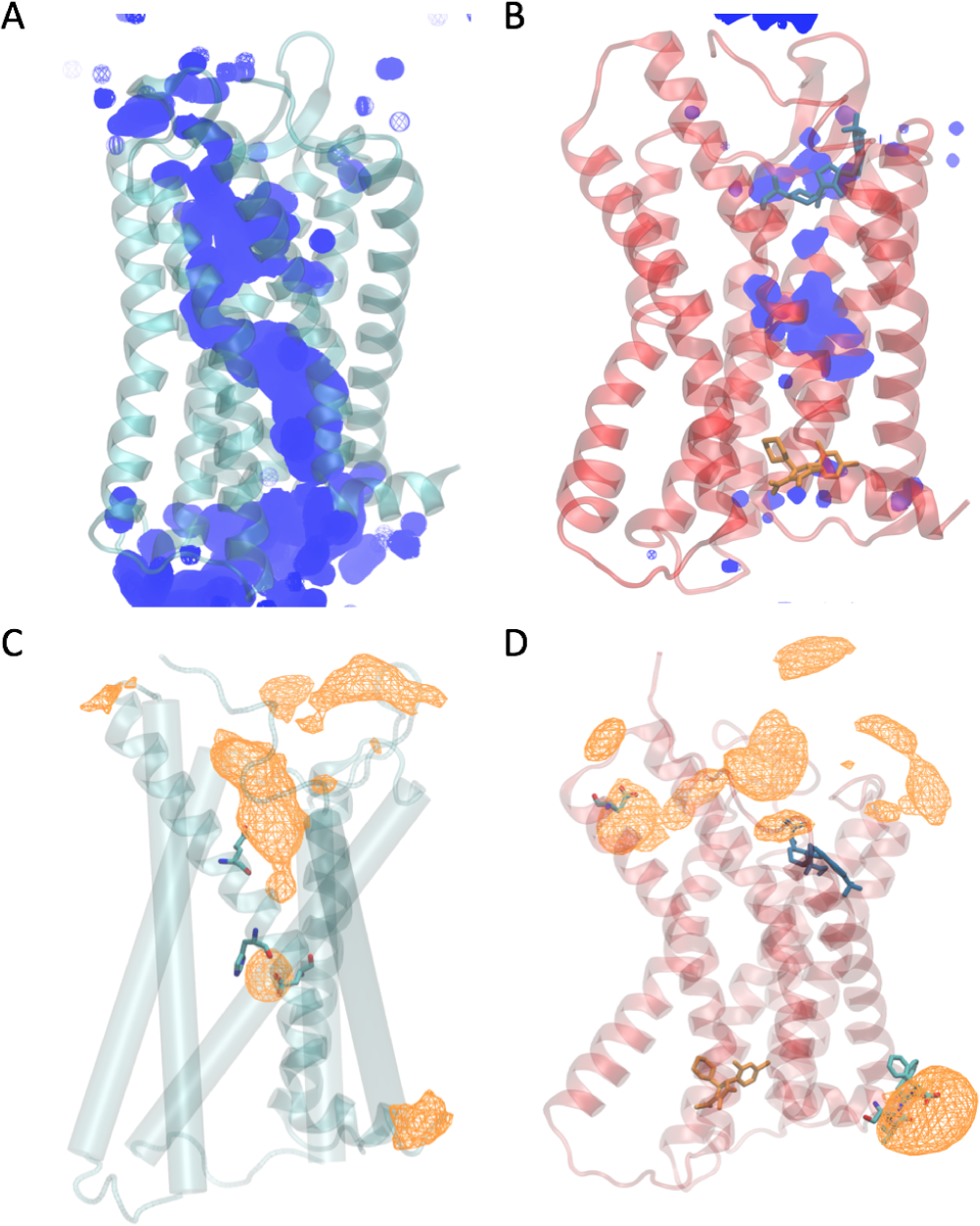
Ligands disrupt a continuous internal water and sodium ion pathway. Average water density over a 50 microsecond simulation of A) Apo (teal) and B) Holo (red). The orthosteric ligand is shown in blue and the allosteric ligand is shown in orange. Total average sodium ion density in C) Apo and D) Holo. Highest occupancy residues depicted in cyan licorice and plotted in Figure S8.

A sodium ion pathway is also disrupted when ligands are bound to CCR2. Sodium ion occupancy per residue was calculated for apo and holo CCR2 (Figure S8B). With ligands bound, sodium ions interact with residues only on the extracellular and intracellular ends of the protein, particularly with residues Glu 270^6.62^ and His 317^8.55^- Lys 320^8.58^. Without ligands, sodium ions interact dramatically more with residues Asp 88^2.50^, Glu 291^7.39^, and His 297^7.45^, which are in the core of the protein. The interaction of Na^+^ ions and Asp 88^2.50^ is thought to be an integral part of receptor activation,^42^ suggesting again that the apo systems are accessing conformations along a pathway toward activation.

## Conclusions

To characterize the basal dynamics of CCR2 and understand how small molecule antagonists modulate these dynamics, we used atomic simulations and MSM theory to compare the metastable states accessed by apo or Holo CCR2. Interestingly, several intermediate states reveal a novel putative binding site that could be targeted with small molecule inhibitors. We found that the kinetics, dynamics, and conformational ensembles of the apo and holo systems differ greatly: antagonists dampen CCR2 dynamics, prevent quick transitions between metastable states, and are associated with different key motions and residue rearrangements. Without antagonists, CCR2 is able to access other distinct metastable states that are likely sampling along an activation pathway. These intermediate states not only inform us about the basal dynamics of CCR2, but may also be useful for rational drug design and modification of previously unsuccessful drugs.

## Materials and Methods

See the Supporting Information for full Materials and Methods.

### System Preparation and Molecular Dynamics Simulations

Two systems were simulated: CCR2 Holo, with both co-crystallized antagonist ligands bound, and CCR2 Apo, without ligands bound. CCR2-RA-[R] and BMS 681^14^ were deleted to build the Apo system. Each all-atom system is embedded in a biologically similar POPC bilayer, explicitly solvated with TIP3P, and simulated with 150mM NaCl, at physiological pH, at 310K and 1 bar. The initial coordinates were taken from the experimental crystal structure^14^ and simulated for 50ns MD simulations on local resources before simulation on Anton2. The Anton2 simulations were run in the NPT ensemble, using Anton’s Berendsen thermostat-barostat, at 310K and 1 bar with a 2-fs time-step and partial mesh Ewald electrostatic approximation. The two systems were simulated for an aggregate total of 260 microseconds.

### Building the Markov State Models

PyEMMA^35^ was used for feature selection, dimensionality reduction, clustering, MSM building, and MSM analysis. Dimensionality reduction using tICA^33,34^ was performed on all trajectories. The Apo and Holo systems were clustered separately using K-means. MSMs were coarse-grained with PCCA+ and transitions were determined with hidden Markov models (HMMs). The models were selected based on implied timescale plots, Chapman-Kolmogorov tests, and visual inspection. Representative states were selected by identifying the most populated cluster in each metastable state, and using MSMBuilder^43^ to find the centroid of that cluster.

## Data Availability

All MD simulation data will be available for download.

## Acknowledgments

The authors thank Tracy Handel and Irina Kufareva for their valuable input and discussions as well as initial modification of CCR2 coordinates. We also thank the organizers and participants of the 2018 Workshop on Free energy Methods, Kinetics and Markov State Models in Drug Design for helpful comments and discussion. This work was supported in part by the Director’s New Innovator Award Program NIH DP2-OD007237, the National Biomedical Computation Resource (NBCR) NIH P41-GM103426, and the National Science Foundation through XSEDE supercomputing resources provided via TG-CHE060073 to R.E.A. C.T.L. also acknowledges support from the NIH Molecular Biophysics Training Program (T32-GM008326) Anton 2 computer time was provided by the Pittsburgh Supercomputing Center (PSC) through Grant R01GM116961 from the National Institutes of Health. The Anton 2 machine at PSC was generously made available by D.E. Shaw Research.

## Author Contributions

B.C.T, C.T.L, and R.E.A. designed research; B.C.T. performed and analyzed research; B.C.T., C.T.L, and R.E.A. wrote the paper.

## References

[1] A. Ben-Baruch. The multifaceted roles of chemokines in malignancy. Cancer and Metastasis Reviews, 25(3):357–371, 2006.

[2] T O’Connor, L Borsig, and M Heikenwalder. CCL2-CCR2 signaling in disease pathogenesis. Endocrine, Metabolic and Immune Disorders - Drug Targets, 15(2):105–118, 2015.

[3] Michelle Solomon, Balaji Balasa, and Nora Sarvetnick. CCR2 and CCR5 chemokine receptors differentially influence the development of autoimmune diabetes in the NOD mouse. Autoimmunity, 43(2):156–163, 2010.

[4] James E Pease and Richard Horuk. Chemokine receptor antagonists: Part 1. Expert Opinion on Therapeutic Patents, 19(1):39–58, jan 2009.

[5] James E Pease and Richard Horuk. Chemokine receptor antagonists: part 2. Expert opinion on therapeutic patents, 19(2):199–221, feb 2009.

[6] D. J. Scholten, M. Canals, D. Mussang, L. Roumen, M.J. Smit, M. Wijtmans, C. de Graaf, H. F. Vischer, and R. Leurs. Pharmacological modulation of chemokine receptor function. Br. J. Pharmacol, 165:1617–1643, 2012.

[7] S.Y. Lim, A.E. Yuzhalin, A.N. Gordon-Weeks, and R.J. Muschel. Targeting the CCL2-CCR2 signaling axis in cancer metastasis. Oncotarget, 7:28697–710, 2016.

[8] R. Solari, J. E. Pease, and M. Begg. Chemokine receptors as therapeutic targets: why arent there more drugs? Eur. J. Pharmacol., 746:363367, 2015.

[9] Richard Horuk. Chemokine receptor antagonists: overcoming developmental hurdles. Nature reviews. Drug discovery, 8(1):23–33, 2009.

[10] https://www.clinicaltrials.gov/, 2018.

[11] Mohsen Shahlaei, Afshin Fassihi, Elena Papaleo, and Morteza Pourfarzam. Molecular dynamics simulation of chemokine receptors in lipid bilayer: a case study on C-C chemokine receptor type 2. Chemical biology & drug design, 82(5):534–545, 2013.

[12] Swapnil Chavan, Shirishkumar Pawar, Rajesh Singh, and M. Elizabeth Sobhia. Binding site characterization of G protein-coupled receptor by alanine-scanning mutagenesis using molecular dynamics and binding free energy approach: application to C-C chemokine receptor-2 (CCR2). Molecular Diversity, 16(2):401–413, 2012.

[13] Gugan Kothandan, Changdev G Gadhe, and Seung Joo Cho. Structural insights from binding poses of CCR2 and CCR5 with clinically important antagonists: a combined in silico study. PloS one, 7(3):e32864, 2012.

[14] Yi Zheng, Ling Qin, Natalia V. Ortiz Zacarías, Henk de Vries, Gye Won Han, Martin Gustavsson, Marta Dabros, Chunxia Zhao, Robert J. Cherney, Percy Carter, Dean Stamos, Ruben Abagyan, Vadim Cherezov, Raymond C. Stevens, Adriaan P. IJzerman, Laura H. Heitman, Andrew Tebben, Irina Kufareva, and Tracy M. Handel. Structure of CC chemokine receptor 2 with orthosteric and allosteric antagonists. Nature, 540(7633):458–461, 2016.

[15] Naomi R. Latorraca, A. J. Venkatakrishnan, and Ron O. Dror. GPCR Dynamics: Structures in Motion. Chemical Reviews, 117(1):139–155, 2017.

[16] A. J. Venkatakrishnan, Xavier Deupi, Guillaume Lebon, Christopher G. Tate, Gebhard F. Schertler, and M. Madan Babu. Molecular signatures of G-protein-coupled receptors. Nature, 494:185–194, 2013.

[17] Qiansen Zhang, Mang Zhou, Lifen Zhao, Hualiang Jiang, and Huaiyu Yang. Dynamic States of the Ligand-Free Class A G Protein-Coupled Receptor Extracellular Side. Biochemistry, 0(0):null, 2018.

[18] V. Katritch, V. Cherezov, and R.C. Stevens. Structure-Function of the G-protein-Coupled Receptor Superfamily. Annual review of pharmacology and toxicology, 53:531–556, 2013.

[19] A. Manglik, T. H. Kim, M. Masureel, C. Altenbach, Z. Y. Yang, D. Hilger, M. T. Lerch, T. S. Kobilka, F. S. Thian, W. L. Hubbell, R. S. Prosser, and B. K. Kobilka. Structural Insights into the Dynamic Process of beta(2)-Adrenergic Receptor Signaling. Cell, 162: 1431 1431, 2015.

[20] V. Katritch, V. Cherezov, and R. C Stevens. Diversity and modularity of G protein-coupled receptor structures. Trends Pharmacol. Sci., 33:17 27, 2012.

[21] R. U. Malik, M. Ritt, B. T. DeVree, R. R. Neubig, R. K. Sunahara, and S. Sivaramakrishnan. Detection of G protein-selective G protein-coupled receptor (GPCR) conformations in live cells. J. Biol. Chem., 288:17167 17178, 2013.

[22] X. J. Yao, G. Velez Ruiz, M. R. Whorton, S. G. F. Rasmussen, B. T. DeVree, X. Deupi, R. K. Sunahara, and B. Kobilka. The effect of ligand efficacy on the formation and stability of a GPCR-G protein complex. Proc. Natl. Acad. Sci. U. S. A., 106:9501 9506, 2009.

[23] R. Nygaard, Y. Z. Zou, R. O. Dror, T. J. Mildorf, D. H. Arlow, A. Manglik, A. C. Pan, C. W. Liu, J. J. Fung, M. P. Bokoch, F. S. Thian, T. S. Kobilka, D. E. Shaw, L. Mueller, R. S. Prosser,, and B. K. Kobilka. The Dynamic Process of beta(2)-Adrenergic Receptor Activation. Cell, 152:532 542, 2013.

[24] S. Bockenhauer, A. Furstenberg, X. J. Yao, B. K. Kobilka, and W. E. Moerner. Conformational dynamics of single G protein-coupled receptors in solution. J. Phys. Chem. B., 115:13328 13338, 2011.

[25] Anton 2: Raising the Bar for Performance and Programmability in a Special-Purpose Molecular Dynamics Supercomputer. In International Conference for High Performance Computing, Networking, Storage and Analysis, SC, number January, pages 41–53, 2014.

[26] Gregory R. Bowman, Vincent a. Voelz, and Vijay S. Pande. Taming the complexity of protein folding. Current Opinion in Structural Biology, 21(1):4–11, 2011.

[27] William C. Swope and Jed W. Pitera. Describing Protein Folding Kinetics by Molecular Dynamics Simulations. J. Phys. Chem. B, 108:6571–6581, 2004.

[28] W.C. Swope, J.W. Pitera, F. Suits, M. Pitman, M. Eleftheriou, B.G. Fitch, R.S. Germain, A. Rayshubski, T.J.C. Ward, Y. Zhestkov, and R. Zhou. Describing Protein Folding Kinetics by Molecular Dynamics Simulations. 2. Example Applications to Alanine Dipeptide and a beta-Hairpin Peptide. J. Phys. Chem. B, 2004.

[29] Nina Singhal, Christopher D. Snow, and Vijay S. Pande. Using path sampling to build better Markovian state models: Predicting the folding rate and mechanism of a tryptophan zipper beta hairpin. Journal of Chemical Physics, 2004.

[30] Robert D. Malmstrom, Christopher T. Lee, Adam T. Van Wart, and Rommie E. Amaro. Application of molecular-dynamics based markov state models to functional proteins. Journal of Chemical Theory and Computation, 2014.

[31] Rommie E. Amaro and Adrian J. Mulholland. Multiscale methods in drug design bridge chemical and biological complexity in the search for cures. Nature Reviews Chemistry, 2: 0148, 2018.

[32] Rommie E. Amaro, Jerome Baudry, John Chodera, Özlem Demir, Andrew McCammon, Yinglong Miao, and Jeremy C. Smith. Ensemble Docking in Drug Discovery. Biophysical Journal, 114:2271–2278, 2018.

[33] C. R. Schwantes and V. S. Pande. Improvements in Markov State Model Construction Reveal Many Non-Native Interactions in the Folding of NTL9. J. Chem. Theory Comput., 9:2000–2009, 2013.

[34] G. Perez-Hernandez, F. Paul, T. Giorgino, G. De Fabritiis, and F Noe. Identification of slow molecular order parameters for Markov model construction. J. Chem. Phys., 139:015102, 2013.

[35] Martin K. Scherer, Benjamin Trendelkamp-Schroer, Fabian Paul, Guillermo Pérez-Hernández, Moritz Hoffmann, Nuria Plattner, Christoph Wehmeyer, Jan Hendrik Prinz, and Frank Noé. PyEMMA 2: A Software Package for Estimation, Validation, and Analysis of Markov Models. Journal of Chemical Theory and Computation, 11(11):5525–5542, 2015.

[36] Jan Hendrik Prinz, Hao Wu, Marco Sarich, Bettina Keller, Martin Senne, Martin Held, John D. Chodera, Christof Schtte, and Frank No?? Markov models of molecular kinetics: Generation and validation. Journal of Chemical Physics, 134(17), 2011.

[37] Theo A Berkhout, Frank E Blaney, Angela M Bridges, David G Cooper, Ian T Forbes, Andrew D Gribble, Pieter H E Groot, Adam Hardy, Robert J Ife, Rejbinder Kaur, Kitty E Moores, Helen Shillito, Jennifer Willetts, and Jason Witherington. CCR2: characterization of the antagonist binding site from a combined receptor modeling/mutagenesis approach. Journal of medicinal chemistry, 46(19):4070–86, 2003.

[38] Spencer E Hall, Allen Mao, Vicky Nicolaidou, Mattea Finelli, Emma L Wise, Belinda Nedjai, Julie Kanjanapangka, Paymann Harirchian, Deborah Chen, Victor Selchau, Sofia Ribeiro, Sabine Schyler, James E Pease, Richard Horuk, and Nagarajan Vaidehi. Elucidation of binding sites of dual antagonists in the human chemokine receptors CCR2 and CCR5. Molecular pharmacology, 75(6):1325–36, 2009.

[39] AD Caliman, SE Swift, Y Wang, Y Miao, and JA. McCammon. Investigation of the conformational dynamics of the apo A2A adenosine receptor. Protein Science: A Publication of the Protein Society, 24:1004–1012, 2015.

[40] D. Kozakov, L.E. Grove, D.R. Hall, T. Bohnuud, S.E. Mottarella, L. Luo, B. Xia, D. Beglov, and S. Vajda. The FTMap family of web servers for determining and characterizing ligand-binding hot spots of proteins. Nature Protocols, 10:733–755, 2015.

[41] S. Yuan, S. Filipek, K. Palczewski, and H. Vogel. Activation of G-protein-coupled receptors correlates with the formation of a continuous internal water pathway. Nature communications, 5:4733, 2014.

[42] Owen N. Vickery, Catarina A. Carvalheda, Saheem A. Zaidi, Andrei V. Pisliakov, Vsevolod Katritch, and Ulrich Zachariae. Intracellular Transfer of Na+ in an Active-State G-Protein-Coupled Receptor. Structure, 26:171–180, 2018.

[43] Kyle A. Beauchamp, Gregory R. Bowman, Thomas J. Lane, Lutz Maibaum, Imran S. Haque, and Vijay S. Pande. MSMBuilder2: Modeling conformational dynamics on the picosecond to millisecond scale. Journal of Chemical Theory and Computation, 7(10):3412–3419, 2011.

